# Serotonin regulates dynamics of cerebellar granule cell activity by modulating tonic inhibition

**DOI:** 10.1101/411017

**Authors:** Elizabeth Fleming, Court Hull

## Abstract

Understanding how afferent information is integrated by cortical structures requires identifying the factors shaping excitation and inhibition within their input layers. The input layer of the cerebellar cortex integrates diverse sensorimotor information to enable learned associations that refine the dynamics of movement. Specifically, mossy fiber afferents relay sensorimotor input into the cerebellum to excite granule cells, whose activity is regulated by inhibitory Golgi cells. To test how this integration can be modulated, we have used an acute brain slice preparation from young adult rats and found that encoding of mossy fiber input in the cerebellar granule cell layer can be regulated by serotonin (5-HT) via a specific action on Golgi cells. We find that 5-HT depolarizes Golgi cells, likely by activating 5-HT2A receptors, but does not directly act on either granule cells or mossy fibers. As a result of Golgi cell depolarization, 5-HT significantly increases tonic inhibition onto both granule cells and Golgi cells. 5-HT-mediated Golgi cell depolarization is not sufficient, however, to alter the probability or timing of mossy fiber-evoked feed-forward inhibition onto granule cells. Together, increased granule cell tonic inhibition paired with normal feed-forward inhibition acts to reduce granule cell spike probability without altering spike timing. These data hence provide a circuit mechanism by which 5-HT can reduce granule cell activity without altering temporal representations of mossy fiber input. Such changes in network integration could enable flexible, state-specific suppression of cerebellar sensorimotor input that should not be learned, or enable reversal learning for unwanted associations.

**New and Noteworthy:** 5-HT regulates synaptic integration at the input stage of cerebellar processing by increasing tonic inhibition of granule cells. This circuit mechanism reduces the probability of granule cell spiking without altering spike timing, thus suppressing cerebellar input without altering its temporal representation in the granule cell layer.

## Introduction

The cerebellum plays a key role in motor learning, particularly by harnessing a wide array of sensorimotor inputs to establish learned associations that modify motor output. The first major site of cerebellar sensorimotor integration is the granule cell layer, where excitatory input carried by mossy fibers diverges onto a vast number of granule cells. This large-scale divergence has long been thought to provide a substrate for pattern separation that distinguishes unique sensorimotor inputs necessary for associative learning. According to classical and recent models, granule cell layer pattern separation is tightly regulated by local inhibition provided by interneurons called Golgi cells (Albus 1971; Billings et al. 2014; Eccles et al. 1967; Marr 1969). As the sole source of synaptic inhibition onto granule cells, Golgi cells can set the excitability of granule cells and hence the transformation of incoming mossy fiber input into granule cell output (Brickley et al. 1996; Chadderton et al. 2004; Duguid et al. 2012; Hamann et al. 2002; Mitchell and Silver 2003). Due to this central role in controlling granule cell excitability, modulation of Golgi cell activity could represent a key mechanism for generating the types of flexible, context-dependent responses that have been observed in the granule cell layer during behavior (Albergaria et al. 2018; Courtemanche et al. 2009; Ozden et al. 2012).

Neuromodulatory inputs to the granule cell layer are well-suited to assert such context-dependent regulation of Golgi cells and cerebellar processing. Specifically, the cerebellum receives significant serotonergic (5-HT) projections from the reticular and raphe nuclei (Bishop 1985; Dieudonne 2001) that are particularly dense in the granule cell layer (Takeuchi 1983). Moreover, 5-HT receptors have been reported to be expressed on both Golgi cells and granule cells, although the latter appears to be restricted to early stages of cerebellar development (Geurts 2002; Oostland et al. 2014). Because 5-HT activation in other brain regions has been associated with flexible reconfiguration and reversal of learned associations (Clarke 2004; Matias et al. 2017), these anatomical observations suggest that cerebellar 5-HT inputs could provide a key mechanism for modulating learning, as well as cerebellar sensorimotor integration.

Here, we demonstrate that 5-HT depolarizes Golgi cells, likely through activation of the 5-HT2A receptor, but does not act directly on either granule cells or mossy fibers in young adult rats. As a result, 5-HT significantly increases tonic inhibition onto both granule cells and Golgi cells. Additionally, although 5-HT produces a net depolarization of Golgi cells that elevates their firing, it does not significantly alter the probability or timing of evoked Golgi cell inhibition onto granule cells. Together, increased tonic inhibition with normal feed-forward inhibition acts to reduce mossy fiber driven spike probability in granule cells without degrading spike timing. These data provide a circuit mechanism by which 5-HT can regulate the reliability of granule cell firing to modify sensorimotor representations. Such changes in network integration could underlie the types of context-dependent gating of sensorimotor input that have been observed in the cerebellum and enable flexible suppression of inputs that should either not be learned or should be unlearned.

## Materials and Methods

### SLICES AND RECORDINGS

All procedures were approved by the Duke University Institutional Animal Care and Use Committee (IACUC), and are in accord with guidelines set by the US National Institutes of Health. Acute sagittal slices (250 µm) were cut from the cerebellar vermis of postnatal day 20-27 male Sprague Dawley rats. Slices were prepared in an ice-cold solution of 130 mM K-gluconate, 15 mM KCl, 0.05 mM EGTA, 20 mM HEPES, and 25 mM glucose (pH 7.4), with 2.5 µM R-CPP. This solution has previously been found to enhance the survival and health of cerebellar Golgi cells (Hull and Regehr 2012; Kanichay and Silver 2008). Slices were then stored in artificial cerebral spinal fluid (aCSF) containing 125 mM NaCl, 26 mM NaHCO_3_, 1.25 mM NaH2PO_4_, 2.5 mM KCl, 1 mM MgCl_2_, 2 mM CaCl_2_, and 25 mM glucose, and equilibrated with 95% O2 and 5% CO2. This solution contains divalent cation concentrations (≤ 3 mM) that permit Golgi cell spontaneous packemaking in acute rat cerebellar slices. Slices were incubated at 34°C for 20 minutes after preparation, then kept at room temperature for up to 6 hours.

Slices were viewed using Dodt Gradient Contrast optics (Scientifica) on an upright microscope (Olympus BX51WI), with a 40x water-immersion objective and a CMOS camera (QImaging, Rolera Bolt). Whole-cell and cell-attached recordings were obtained with patch pipettes (Golgi cells: 3-5 MΩ, granule cells, whole-cell: 6-9 MΩ, granule cells, cell-attached: 10-14 (MΩ) pulled from borosilicate capillary glass (World Precision Instruments) with a Sutter P-1000 micropipette puller. Electrophysiological recordings were performed at 31-33°C.

Spontaneous IPSCs (sIPSCs) were recorded at 0 mV. Evoked excitatory postsynaptic currents (eEPSCs) were recorded at −70 mV. The reversal potential for evoked inhibitory postsynaptic currents (eIPSCs) in granule cells was determined empirically in each experiment by adjusting the membrane potential until no eEPSC was observed. eIPSCs were recorded at a mean holding potential of 3.8 ± 3.3 mV. Voltage-clamp recordings were performed using an internal pipette solution containing: 140 mM Cs-gluconate, 15 mM HEPES, 0.5 mM EGTA, 2 mM TEA-Cl, 2 mM MgATP, 0.3 mM NaGTP, 10 mM phosphocreatine-tris_2_, and 2 mM QX 314-Cl. pH was adjusted to 7.2 with CsOH. Membrane potentials were not corrected for liquid-junction potential.

Current-clamp recordings were performed with an internal solution containing: 150 mM K-gluconate, 3 mM KCl, 10 mM HEPES, 0.5 mM EGTA, 3 mM MgATP, 0.5 mM GTP, 5 Mm phosphocreatine-tris_2_, and 5 mM phosphocreatine-Na_2_. pH was adjusted to 7.2 with KOH. As described previously (Hull and Regehr, 2013), membrane hyperpolarization was avoided before and immediately after break-in to whole-cell mode to prevent immediate firing rate potentiation during Golgi cell recordings. Golgi cells were identified according their size, shape, location and their spontaneous pacemaking, which was verified in cell-attached mode before break-in to whole-cell mode (Hull and Regehr 2012). Golgi cell recordings in which action potential height did not change across the recording were used for afterhyperpolarization analysis.

Current-clamp experiments were performed in synaptic blockers of AMPA receptors (2,3-Dioxo-6-nitro-1,2,3,4-tetrahydrobenzo[f]quinoxaline-7-sulfonamide disodium salt, NBQX, 5 μm), NMDA receptors (3-((*R*)-2-Carboxypiperazin-4-yl)-propyl-1-phosphonic acid, R-CPP, 5 μm), and GABA receptors (6-Imino-3-(4-methoxyphenyl)-1(6*H*)-pyridazinebutanoic acid hydrobromide, SR95531, 5 μm), unless otherwise noted. For experiments blocking voltage-activated sodium channels, tetrodotoxin citrate (Octahydro-12-(hydroxymethyl)-2-imino-5,9:7,10a-dimethano-10a*H*-[1,3]dioxocino[6,5-*d*]pyrimidine-4,7,10,11,12-pentol citrate,TTX,1 µM) was applied to the bath >2 minutes before recording. For all experiments using 5-HT (3-(2-Aminoethyl)-1*H*-indol-5-ol hydrochloride,10 µM), slices were discarded and replaced following each bath application. The 5-HT2A receptor antagonist MDL 100907 ((*R*)-(+)-α-(2,3-Dimethoxyphenyl)-1-[2-(4-fluorophenyl)ethyl]-4-piperinemethanol, 500 nM) was added to the bath >2 minutes prior to or during recordings. No holding current was injected for current clamp experiments. Notably, we find that even when holding current remains stable and negligible in voltage-clamp, granule cells become more excitable in current-clamp in response to injected current within minutes of break-in. Thus, to measure evoked granule cell spiking, we only considered responses to injected current within 2 minutes after break-in. To accommodate this approach, evoked spiking was measured in separate cells for control and 5-HT experiments. For 5-HT evoked granule cell spiking, all measurements were made within 7 minutes of 5-HT application to the bath. All drugs were purchased from Tocris or Abcam.

Electrophysiological data were acquired using a Multiclamp 700B amplifier (Molecular Devices), digitized at 20 kHz with a Digidata 1440A digitizer (Molecular Devices) and low-pass filtered at 10 kHz. Acquisition was controlled using Axon Clampex 10.3 software. Series resistance was monitored in voltage-clamp recordings with a 5 mV hyperpolarizing pulse, and only recordings that remained stable during data collection were used for analysis. An ISO-flex stimulus isolation unit (A.M.P.I.) and glass pipettes (0.25-1 MΩ) filled with aCSF and placed in the white matter tract were used to activate mossy fiber inputs. Stimulating electrodes were placed a minimum distance of approximately 40 µm from recorded cells. Stimulus intensity was adjusted to achieve success rates of approximately 50% for whole-cell recordings of eIPSCs. Disynaptic eIPSCs were identified according to latency (Kanichay and Silver 2008) and sensitivity to NBQX. For cell attached recordings, stimulus intensity was set to achieve success rates of approximately 50% to the second stimulus.

### ANALYSIS AND STATISTICS

Mean action potential (AP) waveforms were generated by averaging 2 s of spikes per condition in each experiment, then averaging these mean waveforms across experiments. Changes in AP waveforms were calculated by subtracting the average waveforms for each individual cell per condition, then by averaging subtracted waveforms across cells. eEPSC and eIPSC latencies were calculated from the time of stimulus onset to the time at 20% of the peak evoked current amplitude. Quantification of average 5-HT-evoked changes in membrane potential, spontaneous inhibition, and evoked currents was performed using the time period of maximal 5-HT effect, as defined by the window where Golgi cell spike rates were maximally elevated (within a 3 minute window, beginning 1 minute after 5-HT application). Synaptic potency, defined as the amplitude of success only trials, was calculated using trials where currents exceeded a threshold of 5 pA below baseline (for EPSCs). Data are presented as mean ± SEM. Data distributions were tested for normality using the Shapiro-Wilk test. For data sets that were non-normally distributed (Figs. 4C and 2D), non-parametric statistical tests were used, with the specific test indicated in the results for each dataset. For normally distributed data, sample means were compared using paired or unpaired, two-tailed Student’s t-tests. Statistics were calculated using Prism 6.0 (Graphpad). Mean values were considered significantly different at p < 0.05. Significant differences are noted by asterisks, with single asterisks representing p < 0.05, two asterisks representing p < 0.01, and three asterisks representing p < 0.001.

## Results

To test whether 5-HT acts presynaptically to modulate excitatory input from mossy fibers entering the granule cell layer or postsynaptically to modulate either of the two principal cell types of the granule cell layer, we performed whole-cell recordings from granule cells and Golgi cells in acute cerebellar slices. First, to determine whether 5-HT can alter granule cell excitability by modulating evoked excitation from mossy fibers, we recorded excitatory post-synaptic currents (EPSCs) in granule cells while stimulating the white matter, before and after applying 5-HT (**Fig. 1**). These experiments revealed no significant effect of 5-HT on evoked EPSC amplitude (**Fig. 1***C*; control = 55.8 ± 12.3, 5-HT = 50.3 ± 12.7 pA, p = 0.75), potency (control = 60.2 ± 12.4, 5-HT = 55.6 ± 13.2 pA, p = 0.79), failure rate (control = 0.15 ± 0.1, 5-HT = 0.20 ± 0.1, p = 0.64) or paired-pulse ratio (control = 0.97 ± 0.05, 5-HT = 0.99 ± 0.08, p = 0.84, n = 10). Next, to test whether 5-HT acts directly on granule cells to modulate their excitability, we performed current-clamp recordings in the presence of synaptic transmission blockers (NBQX 5 µM, R-CPP 5 µM, SR95531 5 µM) (**Fig. 1***D-G*). Following 5-HT (10 µM) bath application, we observed no change in the membrane potential of recorded granule cells (**Fig. 1***E*; control = −75.1 ± 3.0, 5-HT = −75.4 ± 2.7 mV, p = 0.7435, n = 11) and no difference in evoked spiking in response to current injection (**Fig. 1***F, G*; input/output slope: control = 0.302 ± 0.024; 5-HT = 0.301 ± 0.031, p = 0.86). These data suggest that 5-HT does not act on either excitatory glutamatergic mossy fiber inputs or granule cells at the input stage of cerebellar processing.

**Figure 1.**
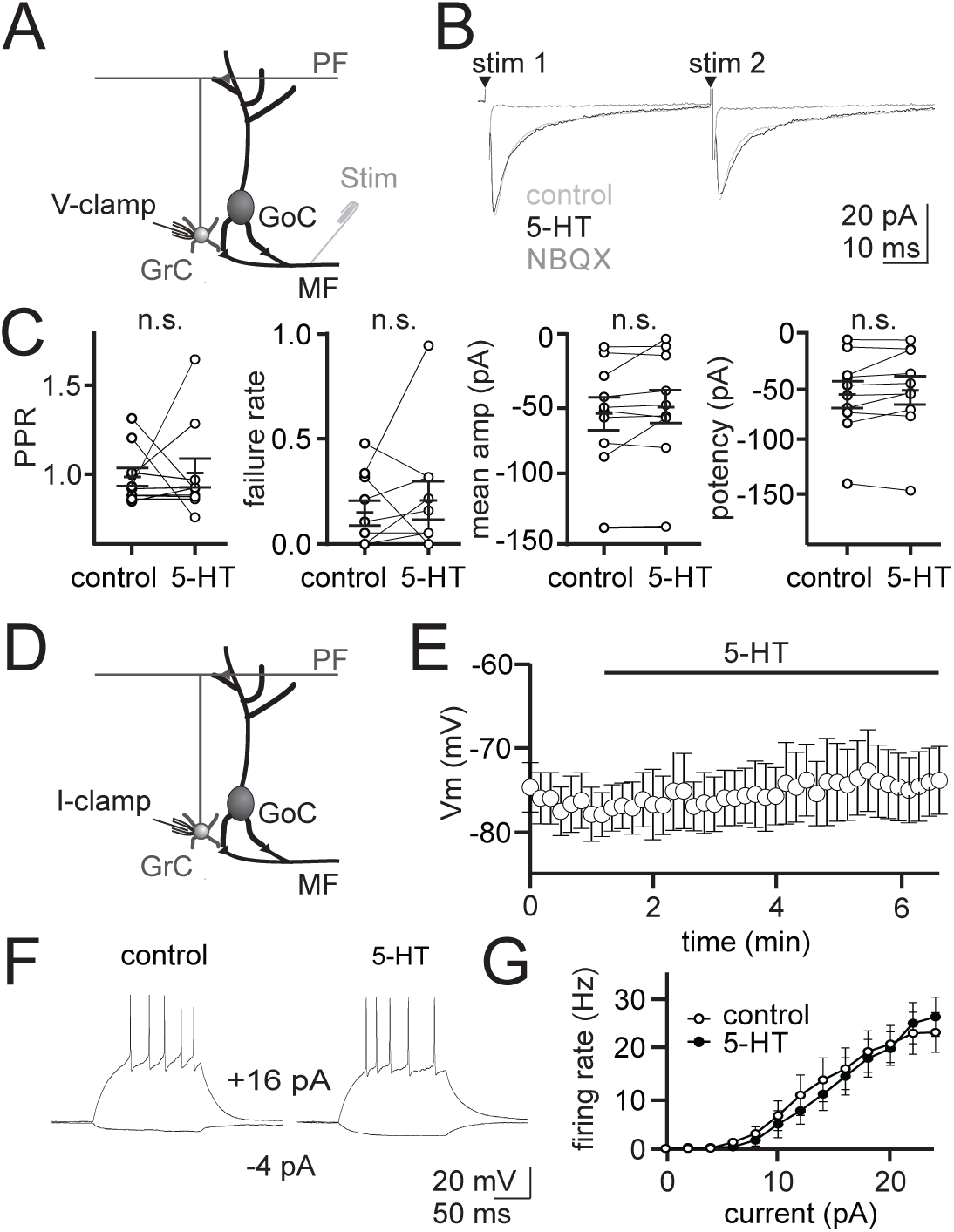
5-HT does not alter evoked excitation onto granule cells or granule cell membrane potential directly. **A.** Schematic of the voltage-clamp recording configuration. **B.** Whole-cell voltage-clamp recordings from an example granule cell. Averaged EPSCs recorded in control, 5-HT (10 μM) and NBQX (5 μM) at - 70 mV. **C.** Summary of paired-pulse ratio, failure rate, mean amplitude, and potency for recorded granule cells. Overall mean and SEM (solid bars) are overlaid (n = 10). n.s. = statistically non-significant. **D.** Schematic of current-clamp recording configuration. **E.** Average membrane potential of granule cells before and after 5-HT application (n = 11) in the presence of NBQX (5 μM) CPP (5 μM) and SR95531 (5 μM). **F.** Whole-cell current-clamp recordings from two example granule cells given a series of 2 pA steps in the presence of NBQX (5 μM) CPP (5 μM) and SR95531 (5 μM). **G.** Average relationship between firing rate and current steps (2 pA steps, input/output relationship) for granule cells with synaptic blockers only (control, n = 13) or synaptic blockers and 5-HT (n = 10).

Next, we tested whether 5-HT can directly modulate the excitability of Golgi cells, the interneurons responsible for all inhibition of cerebellar granule cells. These experiments revealed that 5-HT produced a robust increase in Golgi cell firing rates in the presence of synaptic transmission blockers (**Fig. 2***B-D*; 156.2 ± 54.0% mean increase, control = 4.0 ± 0.9, 5-HT = 7.6 ± 1.1 Hz, p = 0.046, Kolmogorov Smirnov test, n = 10). This increase was followed by a slower return toward baseline despite the continued presence of 5-HT, consistent with the well-documented desensitization of 5-HT receptors (Sullivan Hanley and Hensler 2002). Because immunohistochemistry has suggested 5-HT2A receptors may be expressed by Golgi cells (Geurts 2002), we next tested the possibility that these effects on firing rate were mediated by 5-HT2A receptors using the selective antagonist MDL 100907 (500 nM) (Kehne 1996; Sorenson 1993). Based on our finding that 5-HT-induced firing rate increases can begin returning toward baseline within a few minutes, we conducted these experiments by applying 5-HT in the presence of MDL 100907. In support of the hypothesis that 5-HT2A receptor activation likely mediates spike rate increases, we observed no significant change in Golgi cell firing rates in the presence of MDL 100907 (**Fig. 2***E, F*; Δ spike rate = 0.8 ± 1.0 Hz, n = 4, p = 0.1061). We did, however, observe that Golgi cells were more depolarized in the presence of MDL as compared to control conditions, suggesting a resting serotonergic tone in the slice (**Fig. 2***E*; V_m_ control = −54.8 ± 0.8 mV; V_m_ MDL: −59.1 ± 1.8 mV; p = 0.0328).

**Figure 2.**
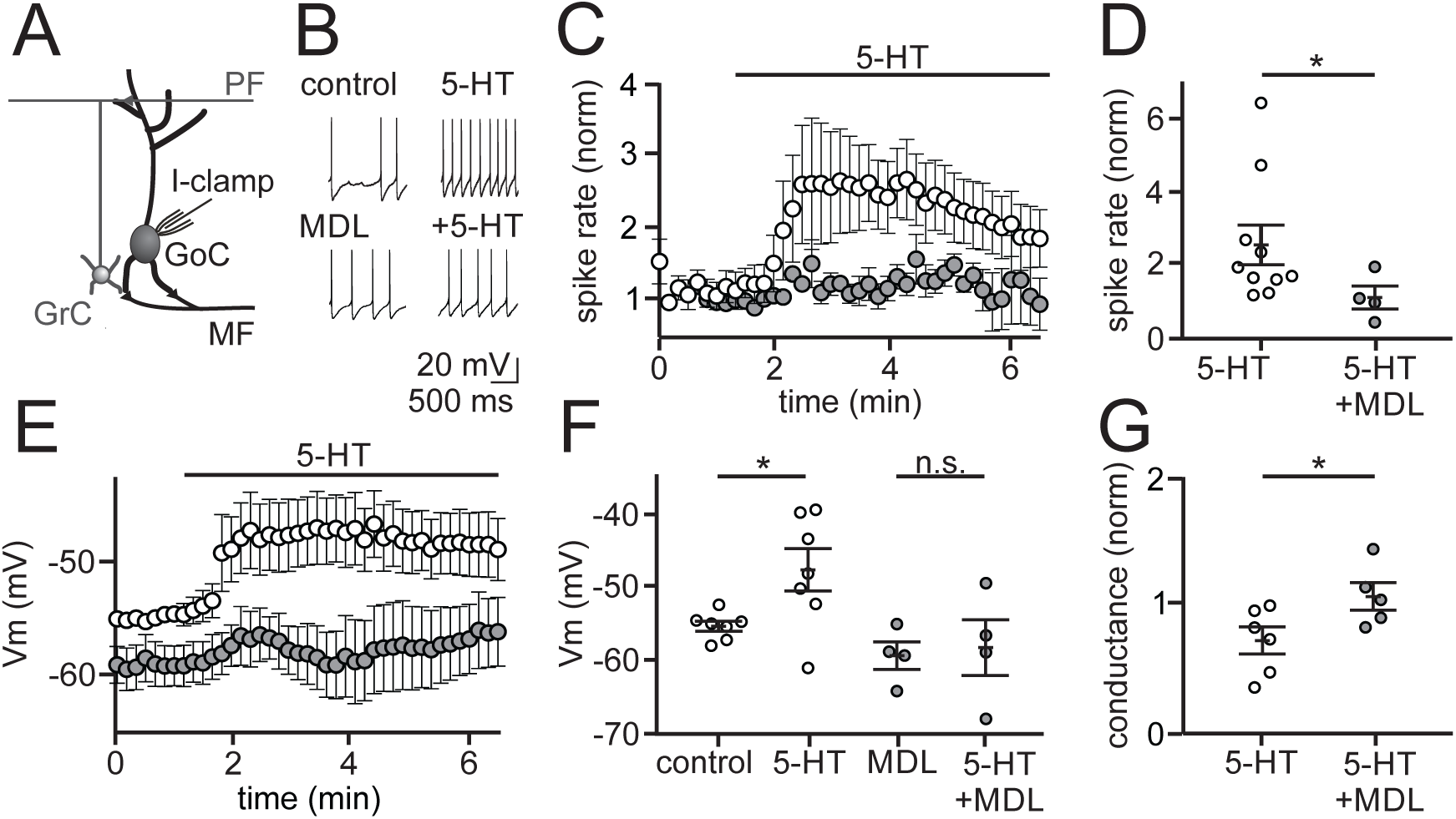
5-HT depolarizes Golgi cells by activating 5-HT2A receptors. **A.** Schematic of the recording configuration. **B.** Example traces from two individual whole-cell, current-clamp recordings from Golgi cells before and during bath application of 5-HT (10 μM), with and without the 5-HT2A receptor-specific antagonist, MDL 100907 (500 nM). NBQX (5 μM) CPP (5 μM) and SR95531 (5 μM) present in all recordings. **C.** Summary of normalized spike rates across time in control (open, n = 10) and in MDL (gray, n = 4). **D.** Summary of mean normalized spike rates during peak 5-HT response window in control and in MDL. **E.** Summary of membrane potential (Vm) over time with 5-HT added to control (open, n = 7) and MDL (gray, n = 4). **F.** Summary of mean membrane potential during control, 5-HT, MDL and MDL + 5-HT. Membrane potentials in 5-HT were measured at the time of peak 5-HT modulation, and all experiments were performed in TTX (1 μM). n.s. = statistically non-significant **G.** Summary of normalized change in conductance during 5-HT application in control (open, n = 6) and MDL (gray, n = 5) as measured by a 5 mV test pulse.

5-HT2A receptor activation has been reported to modulate neuronal excitability by several mechanisms, including directly depolarizing the membrane either by opening a depolarizing conductance (Hannon and Hoyer 2008) or attenuating resting potassium conductance (Villalobos et al. 2005). Thus, we first tested whether 5-HT modulates the resting membrane potential of Golgi cells by recording in current-clamp in the presence of the voltage-activated sodium channel blocker tetrodotoxin (TTX, 1 µM). In these recordings, 5-HT produced a significant depolarization of the Golgi cell membrane (**Fig. 2***E, F*; 7.6 ± 2.4 mV depolarization, control = −55.1 ± 0.7, 5-HT = −47.6 ± 2.9 mV, p = 0.024, n = 7). This depolarization was absent when 5-HT was applied in the presence of MDL 100907 (**Fig. 2***E, F*; 1.1 ± 1.9 mV hyperpolarization, MDL = −58.0 ± 3.8, +5-HT = −59.1 ± 1.9 mV, p = 0.6063, n = 4). Consistent with the hypothesis that this membrane depolarization is mediated by a reduction in resting potassium conductance rather than activation of a depolarizing conductance, we measured an increase in membrane resistance following 5-HT application that did not occur in the presence of MDL (**Fig. 2***G*; 312.9 ± 225.6 MΩ increase, control = 360.6 ± 60.3, 5-HT = 679.7 ± 274.6 MΩ, p = 0.0411, n = 6; MDL: control = 588.8 ± 106.9, 5-HT = 590.5 ± 86.4 MΩ, p = 0.9480, n = 5).

Previous work has demonstrated that Golgi cell spike rate can also be modulated by a plasticity mechanism termed firing rate potentiation (FRP) (Hull et al. 2013). This FRP plasticity mechanism involves a kinase-dependent, bi-directional regulation of BK-type potassium channels participating in the action potential afterhyperpolarization (AHP) (Hull et al. 2013). Specifically, CaMKII activation is thought to increase BK channel open probability, enhancing AHPs and decreasing spike rates, while PKC is thought to decrease BK channel open probability, leading to smaller AHPs and elevated spiking (van Welie and du Lac 2011). Because 5-HT2A receptors act via a Gq-coupled pathway to increase PLC, it is possible that they can also engage this same FRP plasticity mechanism by activating PKC to modulate BK channels and enhance Golgi cell spike rates. To test whether 5-HT can engage FRP in addition to depolarizing Golgi cells, we measured the effect of 5-HT on Golgi cell AHPs in control conditions and after inducing FRP (**Fig. 3**). The rationale for this approach is that, if 5-HT acts in part through an FRP mechanism to modulate BK channels, it should have a smaller effect on AHPs after FRP has already been maximally engaged. Hence, we began by measuring the effects of 5-HT on Golgi cell AHPs in control conditions from a subset of our experiments where there was no change in the amplitude of action potentials across the duration of the experiment. Consistent with previous data showing a small change in the AHP in response to depolarization (Hull et al. 2013), we measured a 1.4 ± 0.2 mV change in the AHP following 5-HT application (**Fig. 3***A*). Next, we compared the change in the Golgi cell AHP induced by a depolarization protocol previously established to produce maximal FRP with the change in the AHP when 5-HT was applied after maximal FRP. Consistent with previous results, FRP induction produced a significant change in the AHP (**Fig. 3***C; D*; Δ = 2.4 ± 0.8 mV, control = 13.6 ± 1.9, FRP = 11.0 ± 1.5 mV, p = 0.0199, n = 4). Following maximal FRP induction, 5-HT produced an additional reduction in the AHP that was not different from the AHP change induced by 5-HT in control conditions (**Fig. 3***C;* 5-HT only = 1.6 ± 0.06, 5-HT with FRP = 1.2 ± 0.2 mV, p = 0.1701, n = 4). Because 5-HT produces the same change in the AHP in control and after FRP induction, we conclude that 5-HT does not act through the previously described FRP mechanism to modulate Golgi cell spike rates.

**Figure 3.**
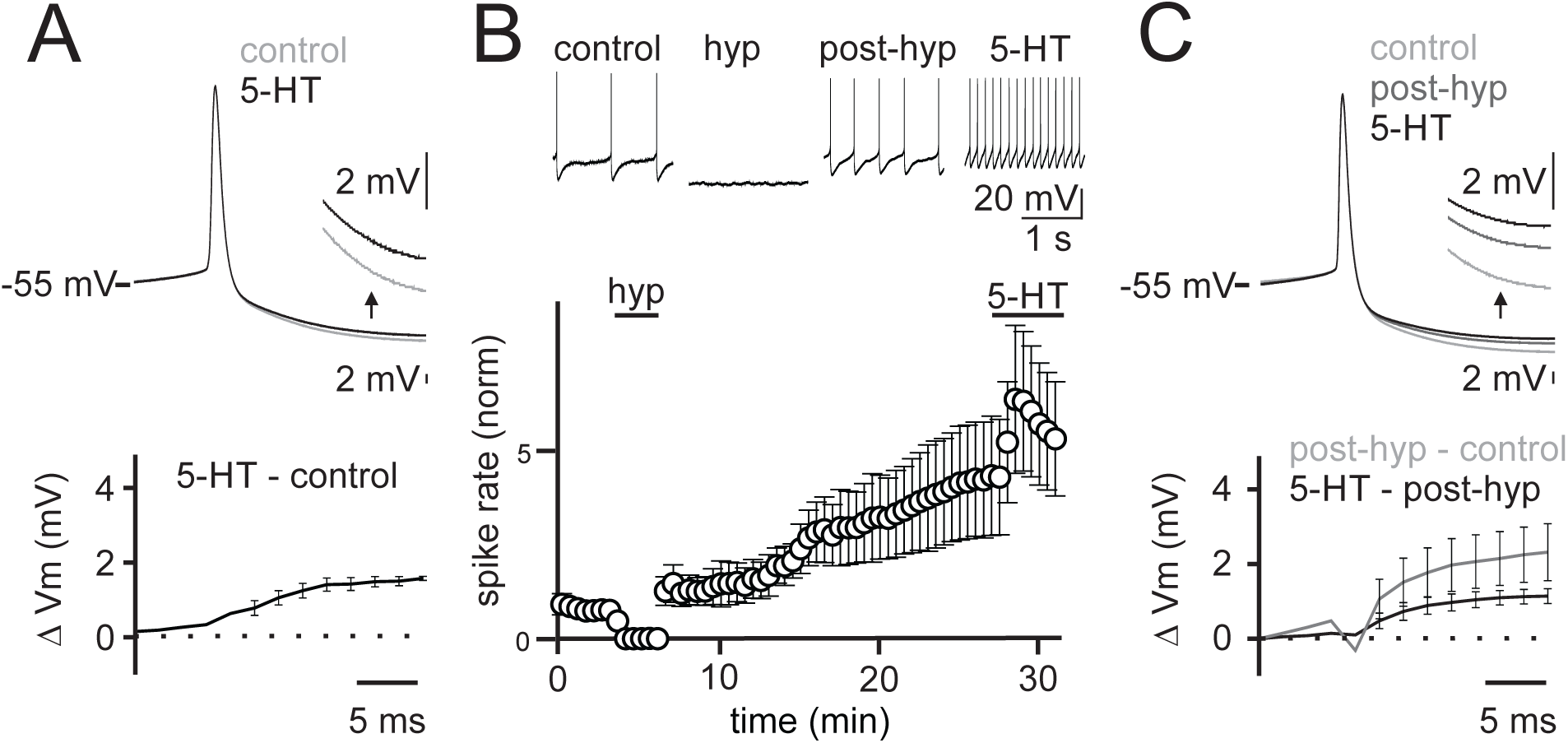
5-HT-mediated increase in Golgi cell excitability is not occluded by hyperpolarization-induced firing rate potentiation. **A.** Top, averaged Golgi cell action potential (AP) waveforms in control and following 5-HT application in the subset of cells with stable AP height (n = 3 cells). Inset shows expanded view of the afterhyperpolarization (AHP). Bottom, subtraction between conditions. NBQX (5 μM) CPP (5 μM) and SR95531 (5 μM) included in all recordings. **B.** Top, Example Golgi cell recording showing action potentials from an experiment where a hyperpolarizing step (−50 pA, 3 min) was applied to induce firing rate potentiation prior to 5-HT application. Bottom, mean normalized firing rate across experiments (n = 5). Error bars are S.E.M. **C.** Top, averaged AP waveforms (n = 4 cells) for each condition. Inset shows expanded view of the afterhyperpolarization (AHP). Bottom, subtractions of AP waveforms between subsequent conditions.

**Figure 4.**
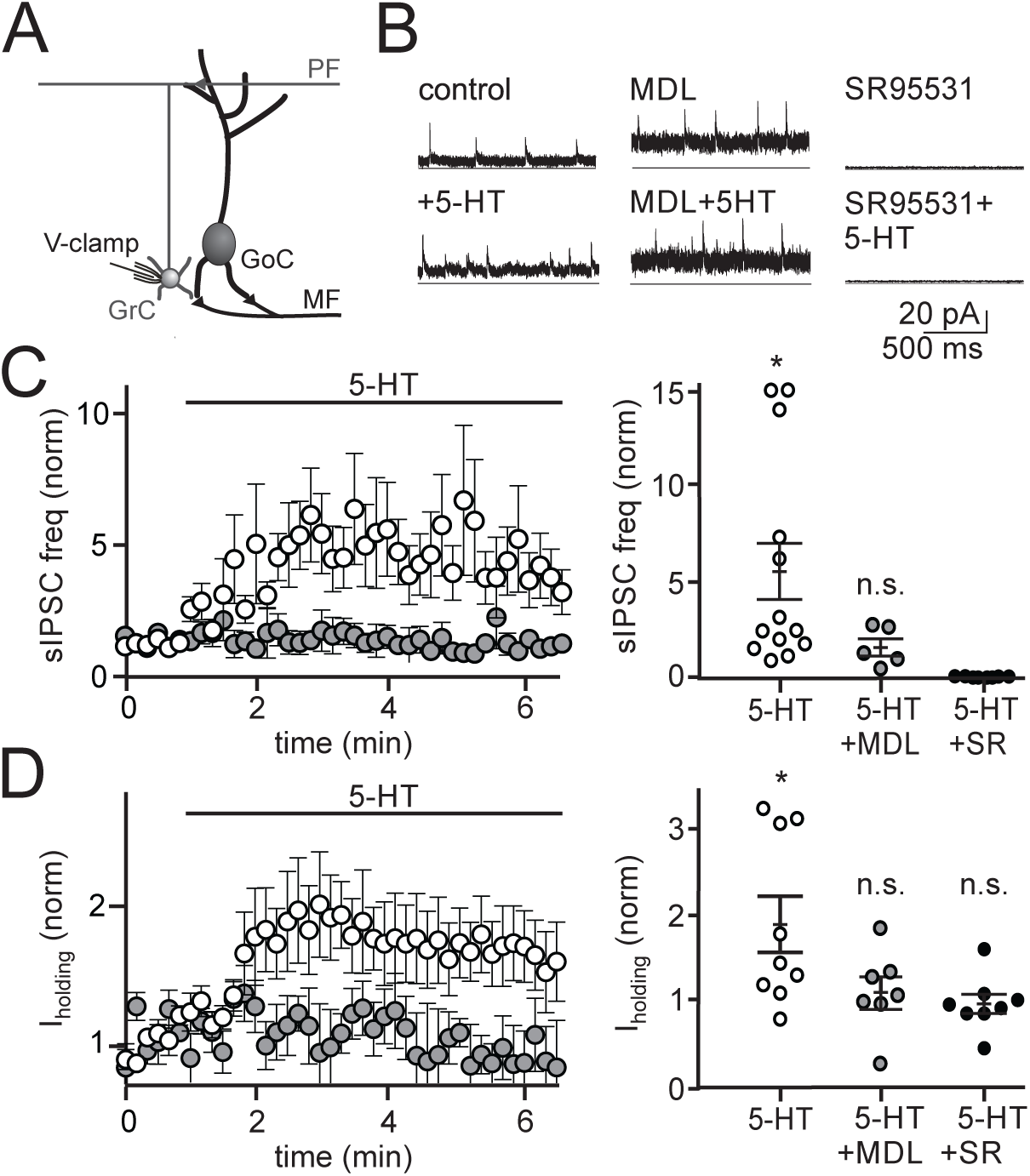
5-HT increases spontaneous inhibition and tonic holding current on granule cells in a 5-HT2AR-dependent manner. **A.** Schematic of the recording configuration. **B.** Example whole-cell, voltage-clamp recordings from 3 granule cells held at the reversal potential for excitation (0mV) before and during bath application of 5-HT (left), with and without the 5-HT2A receptor antagonist MDL 100907 (middle) or SR95531 (right). **C.** Left, Mean normalized spontaneous inhibitory post-synaptic potential (IPSCs) frequency across time in control (open, n = 13) and MDL (gray, n = 5). Right, mean sIPSC frequency quantified at the time of peak 5-HT effect for individual granule cells. Solid black lines are mean and S.E.M. **D.** Left, mean normalized holding current across time in control (open, n = 9) and MDL (gray, n = 7). Right, mean holding current quantified at the time of peak 5-HT effect for individual granule cells. Solid black lines are mean and S.E.M. n.s. = statistically non-significant.

The primary role of Golgi cells is to provide synaptic inhibition to granule cells at the input layer of cerebellar processing. Specifically, GABA release from Golgi cells mediates both tonic and phasic synaptic inhibition of granule cells via distinct types of GABA_A_ receptors (Kaneda et al. 1995; Puia et al. 1994). To test how 5-HT-mediated changes in Golgi cell firing alters each type of granule cell inhibition, we next performed voltage-clamp recordings from granule cells in the absence of synaptic transmission blockers. Following 5-HT bath application, the frequency of spontaneous inhibitory post-synaptic currents (sIPSCs) was significantly increased in recorded granule cells (**Fig. 4***C*; 374.1 ± 148.4% mean increase, control = 2.8 ± 0.9, 5-HT = 9.6 ± 4.0 Hz, p = 0.016, Wilcoxon matched-pairs signed rank test, n = 13), with no associated change in sIPSC amplitude (control = 5.9 ± 0.6 pA, serotonin = 5.7 ± 0.6 pA, p = 0.67, n = 13). In addition, we measured a significant increase in the tonic holding current that reflects GABAergic inhibition via non-desensitizing GABA_A_ receptors (Brickley et al. 1996; Kaneda et al. 1995; Rossi and Hamann 1998; Wall and Usowicz 1997) (**Fig. 4***D*; 83.0 ± 29.1%, control = 8.3 ± 1.3, 5-HT = 13.3 ± 1.8 pA, p = 0.02, n = 9). These large increases in both IPSC frequency and holding current did not occur when 5-HT was applied in the presence of SR5531 (**Fig. 4***C, D*; −2.6 ± 0.1% holding current decrease, SR95531 holding current = 12.5 ± 2.4 pA; SR95531 + 5-HT holding current = 12.0 ± 2.6 pA; p = 0.6308, n=8; no sIPSCs detected in SR95531), or when 5-HT was applied in the presence of MDL 100907 (17.9 ± 26.2% mean sIPSC rate increase, control = 23.7 ± 7.7 Hz, 5-HT = 32.9 ± 16.6 Hz, p = 0.39, n = 5; 21.2 ± 18.8% mean holding current increase, control = 19.6 ± 4.6 pA, 5-HT = 22.3 ± 4.3 pA, p = 0.67, n = 7, respectively). These data suggest that the 5-HT-mediated increases in Golgi cell firing rate can significantly elevate granule cell tonic inhibition.

Golgi cells also make inhibitory synaptic connections onto neighboring Golgi cells (Hull and Regehr 2012), thus 5-HT should also increase inhibition onto Golgi cells. Such a circuit configuration could serve as a form of negative feedback to restrict the magnitude of granule cell inhibition. To test this hypothesis, we next measured how 5-HT modulates synaptic inhibition onto Golgi cells by performing voltage-clamp recordings without blocking synaptic transmission. In these experiments, 5-HT significantly elevated Golgi cell sIPSC frequency (**Fig. 5***C*; 433.9 ± 75.9% mean increase, control = 3.4 ± 0.5, 5-HT = 17.1 ± 1.5 Hz, p = 0.0008, n = 5) consistent with a previous report that had attributed such effects to inputs from Lugaro cells (Dieudonne and Dumoulin 2000). To test the hypothesis that this increase in Golgi cell inhibition serves to regulate the magnitude of 5-HT-induced changes in Golgi cell spontaneous spiking (and hence granule cell spontaneous inhibition), we next performed current-clamp recordings in the absence of synaptic blockers. These experiments revealed that Golgi cell spike rates increased in the presence of 5-HT (**Fig. 5***D*; 79.2 ± 30.5% mean increase, control = 6.2 ± 0.9, 5-HT = 10.8 ± 1.6 Hz, p = 0.081, n = 3), but to a much smaller degree than when synaptic transmission was blocked (156.2 ± 54.0% mean increase, control = 4.0 ± 0.9, 5-HT = 7.6 ± 1.1 Hz, p = 0.046, n = 10). We conclude that the increase in granule cell layer inhibition mediated by 5-HT is restricted by a negative feedback mechanism regulated at least in part by Golgi cell to Golgi cell synapses, and possibly also by additional 5-HT-induced input from Lugaro cells.

**Figure 5.**
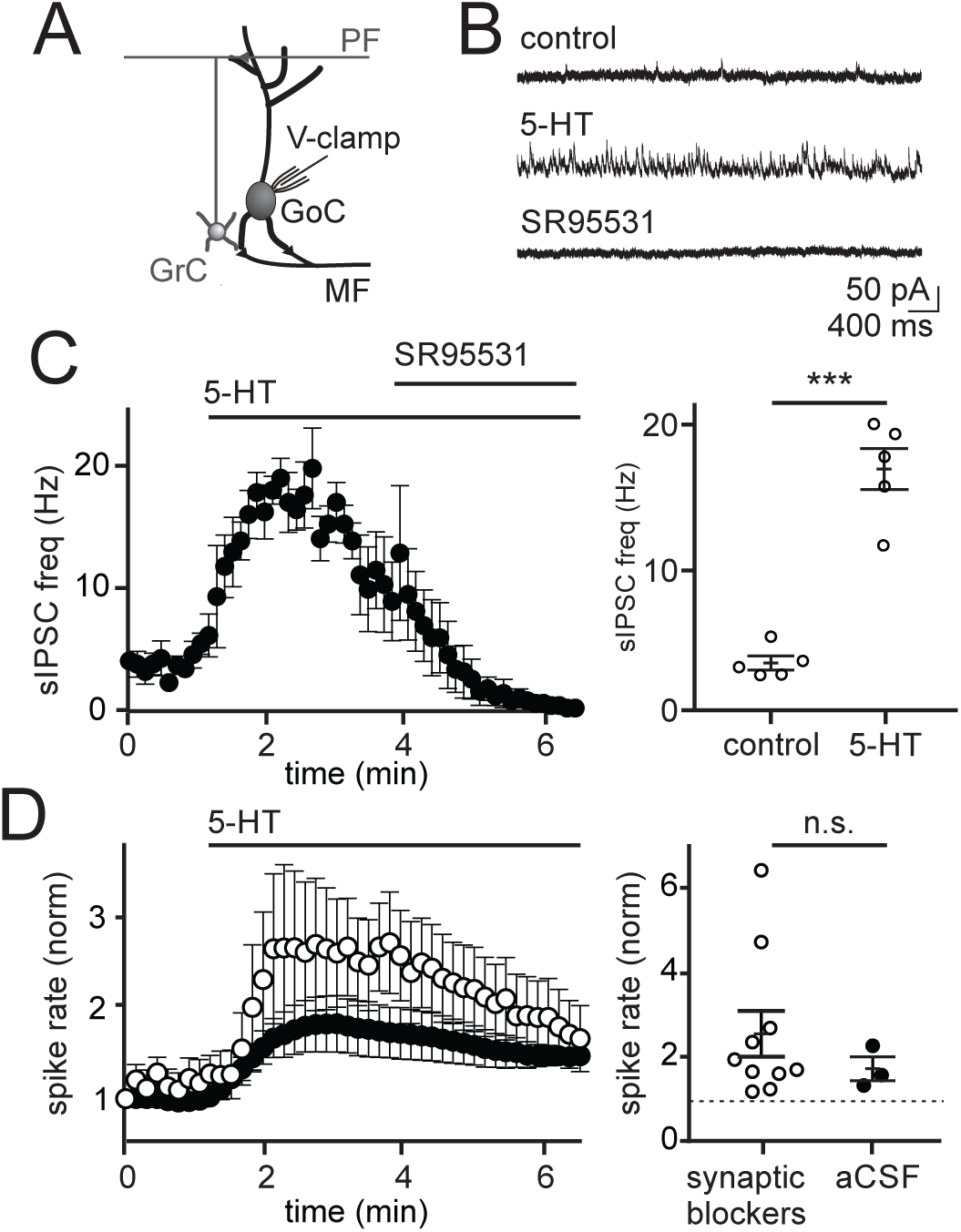
5-HT increases the frequency of sIPSCs on Golgi cells, reducing the magnitude of 5-HT-induced spike rate enhancement. **A.** Schematic of the recording configuration. **B.** Example Golgi cell voltage-clamp recording (0 mV) of IPSCs before and after 5-HT application. Left, mean IPSC frequency before and during application of 5-HT (10 μM; n = 5). Right, mean sIPSC rates measured at baseline and peak response for individual Golgi cells. Solid lines reflect mean and S.E.M. across the population. Left, mean whole-cell, current-clamp Golgi cell spike rate with (open, n = 10) and without (black, n = 3) synaptic transmission blockers (CPP (5 μM), NBQX (5 μM), SR95531 (5 μM)). Right, mean spike rates measured at baseline and peak response for individual Golgi cells. Solid lines reflect mean and S.E.M. across the population. n.s. = statistically non-significant.

In addition to regulating tonic inhibition of granule cells, Golgi cells provide evoked feed-forward and feedback inhibition to granule cells in a manner that is thought be important for regulating spike timing (Kanichay and Silver 2008). Thus, we next tested whether 5-HT acts to regulate evoked (phasic) granule cell inhibition by performing voltage-clamp recordings from granule cells while electrically stimulating mossy fibers (**Fig. 6**). These experiments revealed no statistical differences in the amplitude (**Fig. 6***C, D*; stim 1: control = 19.2 ± 1.7, 5-HT = 17.7 ± 1.9 pA, p = 0.93; stim 2: control = 22.1 ± 2.1, 5-HT = 17.7 ± 1.6 pA, p = 0.48) or latency (stim 1: control = 3.3 ± 0.3, 5-HT = 3.3 ± 0.2 ms, p = 0.66; stim 2: control = 4.0 ± 0.4, 5-HT = 5.0 ± 0.8 ms, p = 0.10, n = 18) of eIPSCs before and after applying 5-HT. Additionally, the failure rate (**Fig. 6***E, F*; stim 1: control = 0.7 ± 0.0, 5-HT = 0.8 ± 0.0, p = 0.30; stim 2: control = 0.8 ± 0.0, 5-HT = 0.8 ± 0.0, p = 0.64) and paired-pulse ratio (control = 1.2 ± 0.1, 5-HT = 1.1 ± 0.1, p = 0.37, n = 18) of eIPSCs were not statistically different before and after applying 5-HT. These data suggest that, while the depolarization evoked by 5-HT is sufficient to increase spontaneous Golgi cell spiking, it is not sufficient to alter the probability or timing of Golgi cell spiking in response to incoming mossy fiber input. Hence, these data reveal that 5-HT acts to regulate spontaneous, tonic granule cell inhibition without significantly altering evoked inhibition.

**Figure 6.**
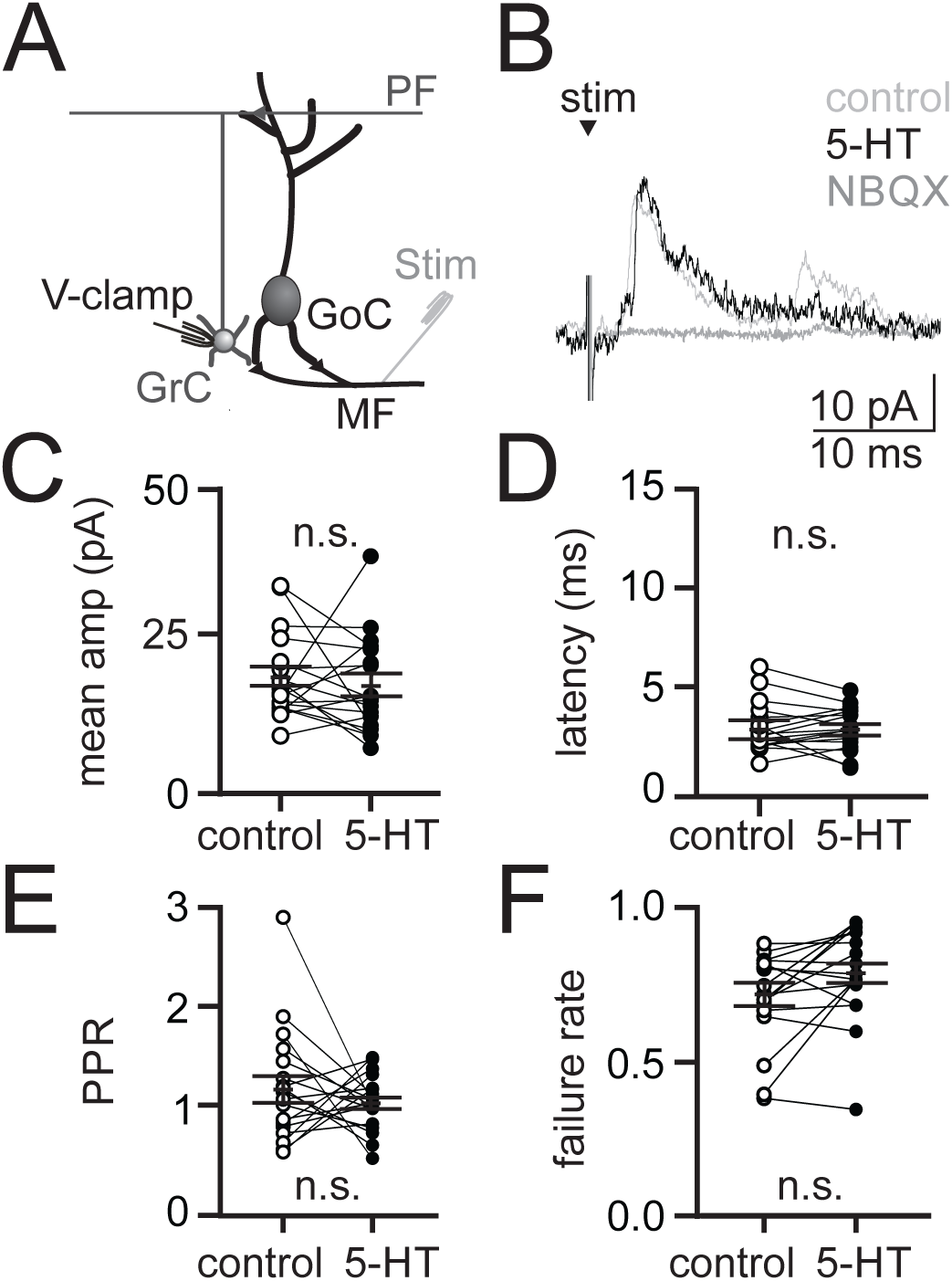
5-HT does not significantly alter evoked feed-forward inhibition onto granule cells. **A.** Schematic of the recording configuration. **B.** Mean evoked IPSCs from an example granule cell held at 0 mV in response to electrical stimulation of the mossy fibers. **C-F.** Mean IPSC amplitude, latency, P P R and failure rate measured from individual granule cells (n = 18). Solid lines reflect mean and S.E.M. across the population. n.s.= statistically non-significant.

Tonic inhibition from Golgi cells regulates granule cell responses to excitatory inputs (Mitchell and Silver 2003), such as those initiated by sensory stimuli (Duguid et al. 2012). Since 5-HT enhances tonic inhibition onto granule cells, we hypothesized that 5-HT may act to selectively reduce efficacy of granule cell responses to mossy fiber input. To test this hypothesis, we performed cell-attached recordings of granule cells while delivering brief bursts of high-frequency stimuli to the mossy fibers to simulate physiological, sensory-evoked cerebellar input (Chadderton et al. 2004) (**Fig. 7**). In most recordings (4 of 6), we observed a marked decrease in the probability of granule cell spiking in the presence of 5-HT (**Fig. 7***D*; stim 1: control = 0.031 ± 0.019, 5-HT = 0.0 ± 0.0, p = 0.10; stim 2: control = 0.075 ± 0.040, 5-HT = 0.038 ± 0.030, p = 0.03; stim 3: control = 0.144 ± 0.019, 5-HT = 0.113 ± 0.022; p = 0.02, n = 4), that quickly recovered above control after application of MDL 100907. The increase in spiking above control levels in MDL is consistent with the finding of a resting serotonergic tone in the slice (**Fig. 2***E*). There was no change in spiking in the remaining 2 cells (*not shown*; stim 1: control = 0.038 ± 0.12, 5-HT = 0.038 ± 0.12, p = 0.5; stim 2: control = 0.075 ± 0.050, 5-HT = 0.075 ± 0.075, p = 0.99; stim 3: control = 0.062 ± 0.062, 5-HT = 0.037 ± 0.037; p = 0.5, n = 2). This result is consistent with the finding that 5-HT did not increase tonic inhibition onto all granule cells (**Fig. 4**). In contrast with spike probability, however, granule cell spike timing was not altered by 5-HT application (**Fig. 7***E*; stim 2: control = 4.0 ± 1.2, 5-HT = 4.5 ± 1.2 ms, p = 0.88; stim 3: control = 4.3 ± 0.60, 5-HT = 4.6 ± 0.60 ms, p = 0.88; n = 4), in agreement with our observations that 5-HT does not alter evoked feed-forward inhibition onto granule cells (**Fig. 6**). These data reveal that 5-HT can act to selectively reduce reliability of granule cell spiking in response to mossy fiber input in a manner that does not alter the timing of granule cell spiking.

**Figure 7.**
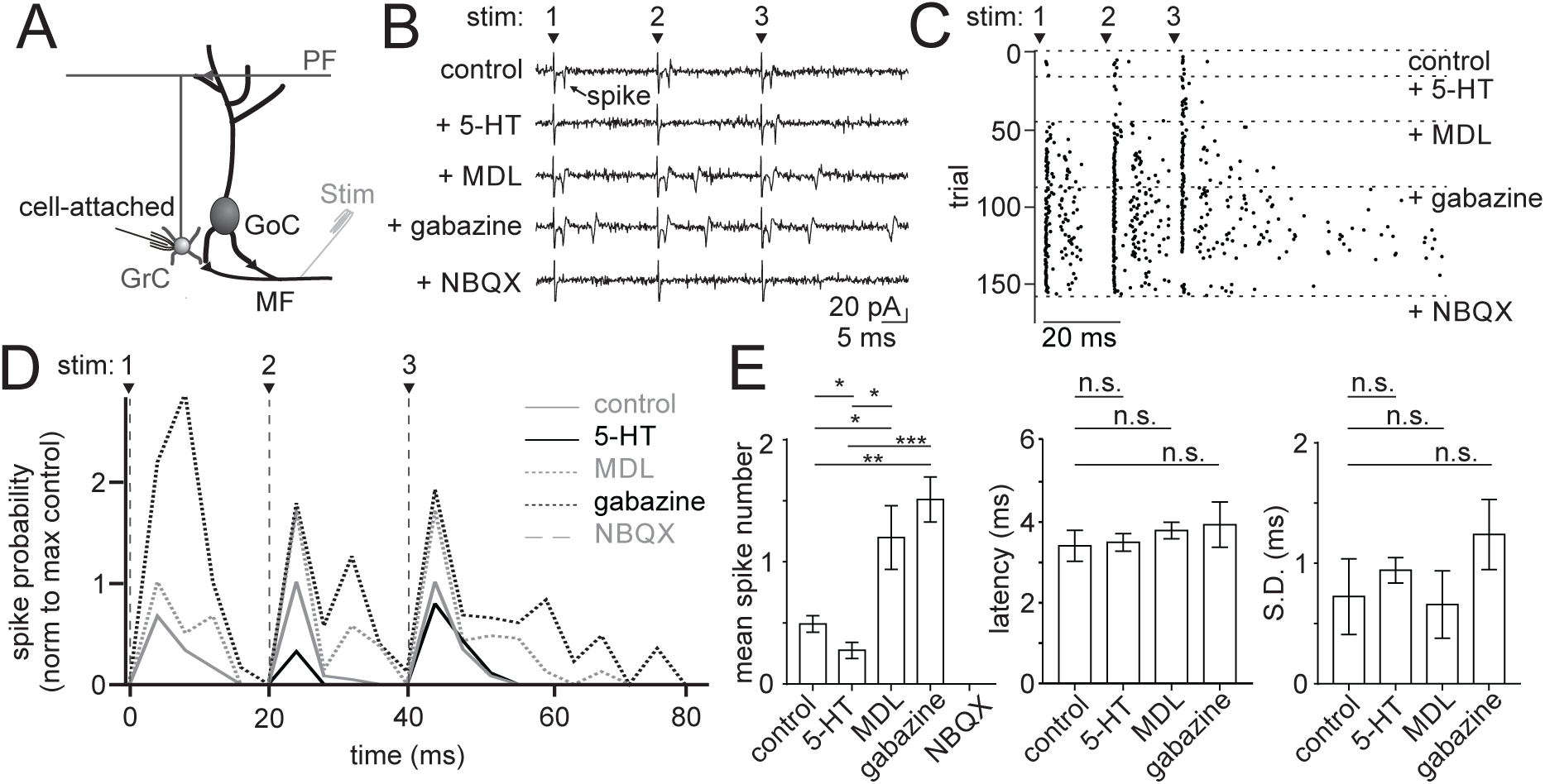
5-HT reduces mossy fiber-evoked spike probability of granule cells in a 5-HT2A-dependent manner. **A.** Schematic of the recording configuration. **B.** Example traces from a cell-attached granule cell recording during electrical stimulation of mossy fibers and sequential bath application of 5-HT, MDL 100907, gabazine and NBQX. **C.** Spike raster from the same example cell in B. **D.** Mean spike probability for granule cells across conditions, (n = 4). **E.** Summary of mean spike number (left), mean first spike latency (middle) and spike jitter (right). n.s. = statistically non-significant.

## Discussion

Here we demonstrate a circuit mechanism by which 5-HT regulates granule cell layer excitability by modulating the activity of cerebellar Golgi cells. By providing a modest depolarization of the Golgi cell membrane that is likely mediated by 5-HT2A receptors, 5-HT acts to selectively modulate the spontaneous spiking of these interneurons without altering their evoked responses to incoming mossy fiber input. As a result, 5-HT acts indirectly to decrease granule cell excitability within a range capable of lowering the probability of granule cell spiking in response to mossy fiber input. However, because 5-HT does not alter evoked inhibition onto granule cells, granule cell spike timing is preserved.

Mechanistically, we find that 5-HT acts to increase inhibition onto both Golgi cells and granule cells by rapidly depolarizing Golgi cells in a cell-autonomous manner, likely through 5-HT2A receptor activation. While the concentration of MDL 100907 we used does not preclude some contribution from other 5-HT receptors, the IC_50_ for the next closest 5-HTR has been reported to be 700 nM (5-HT2C), and all other 5-HT receptors have IC_50_s for MDL that are significantly higher than this (Kehne 1996). Thus, because MDL completely occluded the effects of 5-HT on Golgi cells, it is most likely 5-HT2A receptors are responsible for Golgi cell depolarization. Our results also are consistent with a previous anatomical study that identified 5-HT2A receptors on Golgi cells using histochemical methods (Geurts 2002).

5-HT2A receptors have been suggested to produce neuronal depolarization and increased firing rates via multiple Gq protein-dependent mechanisms, including opening of non-specific cation channels (Hannon and Hoyer 2008) and reducing spike AHPs by decreasing calcium-activated potassium conductances (Villalobos et al. 2005). Our results, however, are not consistent with either of these mechanisms, as 5-HT acts to decrease the Golgi cell membrane conductance and does not alter the spike AHP more than would be predicted by the increased spike rate induced by membrane depolarization (Hull and Regehr 2012). While hyperpolarization of Golgi cells drives a long-term increase in firing rate that results from modification of calcium-activated potassium currents underlying the spike AHP (Hull et al. 2013), our results suggest that the increase in firing rate seen in the presence of 5-HT does not interact with this mechanism.

Instead, the increase in membrane resistance we observe is consistent with reports that 5-HT2A receptors can reduce resting potassium conductances. In particular, these receptors have been shown to reduce an inward-rectifying potassium conductance in cortical fast spiking interneurons, thus depolarizing the membrane potential at rest (Athilingam et al. 2017). 5-HT2A receptors have also been shown to depolarize feed-forward interneurons in the auditory system by reducing a resting potassium conductance (Tang and Trussell 2017). Golgi cells are similar to cortical fast-spiking interneurons, in that they fire narrow action potentials that can achieve high spike rates, they are gap junctionally-coupled within their population, they make a dense, highly divergent axon plexus to release GABA near the soma of their postsynaptic targets, and they provide feed-forward inhibition to their targets. Thus, our data may support the general rule that has been demonstrated in the neocortex and elsewhere that FS-like interneurons commonly express 5-HT2A receptors (Jakab 1998; Kruglikov and Rudy 2008).

Notably, our results differ from an earlier study that suggested no direct effect of 5-HT on Golgi cells in acute slices from the rat cerebellum (Dieudonne and Dumoulin 2000). Though it is unclear why this study did not reveal any effect of 5-HT on Golgi cell spiking, we note that both studies identified increases in synaptic inhibition onto Golgi cells in response to 5-HT. Our results are, however, consistent with the circuit arrangement predicted from the more recent finding that Golgi cells can inhibit one another (Hull et al. 2013). Moreover, our finding that 5-HT reliably increases inhibition onto granule cells is consistent with the depolarizing effect of 5-HT on Golgi cells, as these are the only interneurons in the cerebellar cortex known to inhibit granule cells. In addition, our results are not in conflict with the previously demonstrated result that 5-HT can depolarize Lugaro cells (Dieudonne and Dumoulin 2000). While a physiological connection from Lugaro cells to Golgi cells remains to be demonstrated, inhibition from Lugaro cells in the presence of 5-HT could also contribute to the increases in Golgi cell spontaneous inhibition we observe.

We also note that our results differ from a previous study that showed no effect of 5-HT on granule cell tonic inhibition (Rossi et al. 2003). This study used recording solutions that, in our experience, do not promote Golgi cell spontaneous spiking (methods). Thus, it is possible that conditions in this previous study were not specifically tailored for revealing an effect of 5-HT on granule cell spontaneous inhibition. We therefore speculate that methodological differences may account for the differences between our work and this study.

At the circuit level, it is somewhat surprising that our cell-attached recordings revealed no significant change in the timing of evoked granule cell spiking in 5-HT, even though the time to drive granule cells to spike threshold should be prolonged when inhibition is enhanced. However, we suggest that the rapid arrival of feed-forward inhibition, which remains intact in 5-HT, maintains granule cell spike timing (Duguid et al. 2015; Kanichay and Silver 2008). Specifically, while early spikes remain possible in response to large synaptic inputs, feed-forward inhibition prevents late spikes from occurring. As a consequence, we speculate that the lower spike probability in 5-HT results at least in part from mossy fiber input that cannot drive the granule cells to spike threshold before the arrival of feed-forward inhibition. In support of this view, the time window for granule cell spike generation is dramatically enhanced when feed-forward (and spontaneous) inhibition is blocked with SR95531. Moreover, while it might be expected that early spikes should arrive even sooner due a decrease in the membrane time constant when tonic inhibition is elevated, granule cells have unusually high membrane resistance (<1 GΩ) and small membrane capacitance (∼ 3 pF) and, thus, a rapid membrane time constant (∼5-10 ms). As a result of these passive properties, elevating tonic inhibition at levels measured here (with resistance decreases of approximately 70 mΩ) should only decrease the membrane time constant by hundreds of microseconds.

Granule cell spike timing is thought to be an important feature of sensorimotor encoding at the input stage of cerebellar processing (D’Angelo and De Zeeuw 2009; Kennedy et al. 2014). Recent work has demonstrated that granule cells can receive multimodal input from distinct mossy fibers with different strengths and short-term plasticity (Chabrol et al. 2015). As a result, individual granule cells exhibit diverse spike patterns in response to unique patterns of mossy fiber input that are best distinguished by their timing. Accordingly, it is thought that temporal coding, along with population identity, provides a key mechanism for establishing sensorimotor representations and allowing pattern separation in the granule cell layer (Medina et al. 2000). In this context, our results are consistent with a model in which 5-HT can reduce the strength of a given sensorimotor representation by reducing the reliability of granule cell spiking without altering identity of that representation by preserving temporal coding. Moreover, the precise timing and reliability of granule cell inputs to Purkinje cells determines their potential for undergoing long-term synaptic plasticity (D’Angelo and De Zeeuw 2009). Thus, by selectively reducing the reliability of granule cell spiking, we speculate that 5-HT can decrease associative learning for sensorimotor inputs that behavioral context dictates should not be learned by downstream Purkinje cells. Such a mechanism could also promote reversal learning for an association that has already been made.

Indeed, 5-HT signaling in other brain regions has been associated with reversal learning and behavioral changes that are required for adaptation to new environmental conditions. Specifically, both Dorsal Raphe activation and endogenous 5-HT are necessary for some forms of reversal learning, with disruptions in either resulting in perseverative errors (Clarke 2004; Matias et al. 2017). Moreover, Dorsal Raphe neurons respond to both positive and negative prediction errors, as well as aversive stimuli and changes in motosensory gain (Cohen et al. 2015; Kawashima et al. 2016; Matias et al. 2017), suggesting that 5-HT may serve as a broad signal for behavioral adaption. 5-HT has also been shown to regulate sensory processing, notably by both suppressing auditory inputs and enhancing multisensory inputs in the cerebellar-like dorsal cochlear nucleus (Tang and Trussell 2017; 2015).

The cerebellum receives its serotonergic input from reticular and raphe nuclei (Bishop 1985; Dieudonne 2001), regions that also innervate several other regions of the brain. Thus, 5-HT may be released in the granule cell layer during the same conditions and affect similar learning goals as have been described elsewhere. Given the extensive variety of sensory, motor, and cognitive contextual information converging on the granule cell layer, such adaptive 5-HT signals are poised to influence motor output in response to environmental changes. Hence, by regulating granule cell layer excitability, the circuit mechanisms described here may allow 5-HT to regulate patterns of granule cell activity according to behavioral context in order to regulate motor learning and motor output.

## Author Contributions

EF and CH designed the study, conducted experiments, and wrote the manuscript.

## Acknowledgements

This work was supported by grants from the NIH NINDS (5R01NS096289-02), the Sloan Foundation, and the Whitehall Foundation. We thank Dr. Lindsey Glickfeld and members of the Hull and Glickfeld labs for input and technical assistance throughout the project.

## References

Albergaria C, Silva NT, Pritchett DL, and Carey MR. Locomotor activity modulates associative learning in mouse cerebellum. Nat Neurosci 21: 725–735, 2018.

Albus JS. A theory of cerebellar function. Mathematical Biosciences 10: 25–61, 1971.

Athilingam JC, Ben-Shalom R, Keeshen CM, Sohal VS, and Bender KJ. Serotonin enhances excitability and gamma frequency temporal integration in mouse prefrontal fast-spiking interneurons. Elife 6: 2017.

Billings G, Piasini E, Lorincz A, Nusser Z, and Silver RA. Network Structure within the Cerebellar Input Layer Enables Lossless Sparse Encoding. Neuron 83: 960–974, 2014.

Bishop GA, Ho, R.H. The distribution and origin of serotonin immunoreactivity in the rat cerebellum. Brain Res 331: 195–207, 1985.

Brickley SG, Cull-Candy SG, and Farrant M. Development of a tonic form of synaptic inhibition in rat cerebellar granule cells resulting from persistent activation of GABAA receptors. J Physiol 497 (Pt 3): 753–759, 1996.

Chabrol FP, Arenz A, Wiechert MT, Margrie TW, and DiGregorio DA. Synaptic diversity enables temporal coding of coincident multisensory inputs in single neurons. Nat Neurosci 18: 718–727, 2015.

Chadderton P, Margrie TW, and Hausser M. Integration of quanta in cerebellar granule cells during sensory processing. Nature 428: 856–860, 2004.

Clarke HF, Dalley, J.W., Crofts, H.S., Robbins, T.W., Roberts, A.C. Cognitive inflexibility after prefrontal serotonin depletion. Science 304: 878–880, 2004.

Cohen JY, Amoroso MW, and Uchida N. Serotonergic neurons signal reward and punishment on multiple timescales. Elife 4: e06346, 2015.

Courtemanche R, Chabaud P, and Lamarre Y. Synchronization in primate cerebellar granule cell layer local field potentials: basic anisotropy and dynamic changes during active expectancy. Front Cell Neurosci 3: 6, 2009.

D’Angelo E, and De Zeeuw CI. Timing and plasticity in the cerebellum: focus on the granular layer. Trends in neurosciences 32: 30–40, 2009.

Dieudonne S. Serotonergic Neuromodulation in the Cerebellar Cortex: Cellular, Synaptic, and Molecular Basis. Neuroscientist 7: 207–219, 2001.

Dieudonne S, and Dumoulin A. Serotonin-driven long-range inhibitory connections in the cerebellar cortex. J Neurosci 20: 1837–1848, 2000.

Duguid I, Branco T, Chadderton P, Arlt C, Powell K, and Hausser M. Control of cerebellar granule cell output by sensory-evoked Golgi cell inhibition. Proc Natl Acad Sci U S A 112: 13099–13104, 2015.

Duguid I, Branco T, London M, Chadderton P, and Hausser M. Tonic inhibition enhances fidelity of sensory information transmission in the cerebellar cortex. J Neurosci 32: 11132–11143, 2012.

Eccles J, Ito M, and Szentágothai J. The Cerebellum as a Neuronal Machine. Berlin: Springer, 1967, p. 335–335.

Geurts FJ, De Shutter, E., Timmermans, J. Localization of 5-HT2A, 5-HT3, 5-HT5A and 5-HT7 receptor-like immunoreactivity in the rat cerebellum. J Chem Neuroanat 24: 65–74, 2002.

Hamann M, Rossi DJ, and Attwell D. Tonic and spillover inhibition of granule cells control information flow through cerebellar cortex. Neuron 33: 625–633, 2002.

Hannon J, and Hoyer D. Molecular biology of 5-HT receptors. Behav Brain Res 195: 198–213, 2008.

Hull C, and Regehr WG. Identification of an inhibitory circuit that regulates cerebellar Golgi cell activity. Neuron 73: 149–158, 2012.

Hull CA, Chu Y, Thanawala M, and Regehr WG. Hyperpolarization induces a long-term increase in the spontaneous firing rate of cerebellar Golgi cells. J Neurosci 33: 5895–5902, 2013.

Jakab RL, Goldman-Rakic, P.S. 5-hydroxytryptamine2A sertonin receptors in the primate cerebral cortex: possible site of action of hallucinogenic and antipsychotic drugs in pyramidal cell apical dendrites. Proc Natl Acad Sci U S A 95: 735–740, 1998.

Kaneda M, Farrant M, and Cull-Candy SG. Whole-cell and single-channel currents activated by GABA and glycine in granule cells of the rat cerebellum. J Physiol 485: 419–435, 1995.

Kanichay RT, and Silver RA. Synaptic and cellular properties of the feedforward inhibitory circuit within the input layer of the cerebellar cortex. J Neurosci 28: 8955–8967, 2008.

Kawashima T, Zwart MF, Yang CT, Mensh BD, and Ahrens MB. The Serotonergic System Tracks the Outcomes of Actions to Mediate Short-Term Motor Learning. Cell 167: 933–946 e920, 2016.

Kehne JH, Baron, B.M., Carr, A.A., Shaney, S.F., Elands, J., Feldman, D.J., Frank, R.A., Van Giersbergen, P.L.M., McCloskey, T.C., Johnson, M.P., McCarty, D.R., Poirot, M., Senyah, Y., Siegel, B.W., Widmaier, C. Preclinical characterization of the potential of the putative atypical antipsychotic MDL 100,907 as the potent 5-HT2A antagonist with a favorable CNS safety profile. The Journal of pharmacology and experimental therapeutics 277: 968–981, 1996.

Kennedy A, Wayne G, Kaifosh P, Alvina K, Abbott LF, and Sawtell NB. A temporal basis for predicting the sensory consequences of motor commands in an electric fish. Nat Neurosci 17: 416–422, 2014.

Kruglikov I, and Rudy B. Perisomatic GABA release and thalamocortical integration onto neocortical excitatory cells are regulated by neuromodulators. Neuron 58: 911–924, 2008.

Marr D. A theory of cerebellar cortex. J Physiol 202: 437–470, 1969.

Matias S, Lottem E, Dugue GP, and Mainen ZF. Activity patterns of serotonin neurons underlying cognitive flexibility. Elife 6: 2017.

Medina JF, Garcia KS, Nores WL, Taylor NM, and Mauk MD. Timing mechanisms in the cerebellum: testing predictions of a large-scale computer simulation. The Journal of neuroscience: the official journal of the Society for Neuroscience 20: 5516–5525, 2000.

Mitchell SJ, and Silver RA. Shunting inhibition modulates neuronal gain during synaptic excitation. Neuron 38: 433–445, 2003.

Oostland M, Buijink MR, Teunisse GM, von Oerthel L, Smidt MP, and van Hooft JA. Distinct temporal expression of 5-HT(1A) and 5-HT(2A) receptors on cerebellar granule cells in mice. Cerebellum 13: 491-500, 2014.

Ozden I, Dombeck DA, Hoogland TM, Tank DW, and Wang SS. Widespread state-dependent shifts in cerebellar activity in locomoting mice. PLoS One 7: e42650, 2012.

Puia G, Costa E, and Vicini S. Functional diversity of GABA-activated Cl-currents in Purkinje versus granule neurons in rat cerebellar slices. Neuron 12: 117–126, 1994.

Rossi DJ, and Hamann M. Spillover-mediated transmission at inhibitory synapses promoted by high affinity alpha6 subunit GABA(A) receptors and glomerular geometry. Neuron 20: 783–795, 1998.

Rossi DJ, Hamann M, and Attwell D. Multiple modes of GABAergic inhibition of rat cerebellar granule cells. J Physiol 548: 97–110, 2003.

Sorenson SM, Kehne, J.H., Fadayel, G.M., Humphreys, T.M., Ketteler, H.J., Sullivan, C.K., Taylor, V.L., Schmidt, C.J. Characterization of the 5-HT2 receptor antagonist MDL 100907 as a putative atypical antipsychotic: behavioral, electrophysiological and neurochemical studies. The Journal of pharmacology and experimental therapeutics 266: 684–691, 1993.

Sullivan Hanley NR, and Hensler JG. Mechanisms of ligand-induced desensitization of the 5-hydroxytryptamine2A receptor. The Journal of pharmacology and experimental therapeutics 300: 468-477, 2002.

Takeuchi Y, Kimura, H., Matsuura, T., Yonezawa, T., Sano, Y. Distribution of serotonergic neurons in the central nervous system: a peroxidase-antiperoxidase study with anti-serotonin antibodies. J Histochem Cytochem 31: 181–185, 1983.

Tang ZQ, and Trussell LO. Serotonergic Modulation of Sensory Representation in a Central Multisensory Circuit Is Pathway Specific. Cell Rep 20: 1844–1854, 2017.

Tang ZQ, and Trussell LO. Serotonergic regulation of excitability of principal cells of the dorsal cochlear nucleus. J Neurosci 35: 4540–4551, 2015.

van Welie I, and du Lac S. Bidirectional control of BK channel open probability by CAMKII and PKC in medial vestibular nucleus neurons. J Neurophysiol 105: 1651–1659, 2011.

Villalobos C, Beique JC, Gingrich JA, and Andrade R. Serotonergic regulation of calcium-activated potassium currents in rodent prefrontal cortex. Eur J Neurosci 22: 1120–1126, 2005.

Wall MJ, and Usowicz MM. Development of action potential-dependent and independent spontaneous GABAA receptor-mediated currents in granule cells of postnatal rat cerebellum. Eur J Neurosci 9: 533–548, 1997.

